# A Drosophila Toolkit for Imaging of HA-tagged Proteins Unveiled a Block in Autophagy Flux in the Last Instar Larval Fat Body

**DOI:** 10.1101/2021.05.18.444637

**Authors:** Tadayoshi Murakawa, Tsuyoshi Nakamura, Kohei Kawaguchi, Futoshi Murayama, Zhao Ning, Timothy J Stasevich, Hiroshi Kimura, Naonobu Fujita

## Abstract

For *in vivo* functional analysis of a protein of interest (POI), multiple transgenic strains with POI harboring different tags are needed but generation of these strains is still labor-intensive work. To overcome this, we developed a versatile Drosophila toolkit with a genetically encoded single-chain variable fragment for the HA epitope tag: “HA Frankenbody”. This system allows various analyses of HA-tagged POI in live tissues by simply crossing an HA Frankenbody fly with an HA-tagged POI fly. Strikingly, the GFP-mCherry tandem fluorescent-tagged HA Frankenbody revealed a block in autophagic flux and an accumulation of enlarged autolysosomes in the last instar larval and prepupal fat body. Autophagy was dispensable for the swelling of lysosomes, indicating that lysosomal activity is downregulated at this stage. Furthermore, forced activation of lysosomes by fat body-targeted overexpression of Mitf, the single MiTF/TFE family gene in Drosophila, suppressed the lysosomal swelling and resulted in pupal lethality. Collectively, we propose that downregulated lysosomal function in the fat body plays a role in the metamorphosis of Drosophila.

## Introduction

Protein functional analysis is fundamental in cell and developmental biology. Multiple transgenic strains with a protein of interest (POI) harboring different tags are needed for *in vivo* functional analysis(Kanca et al., 2017). For example, a transgenic line with an epitope tag-fused construct for immunostaining or immunoprecipitation (Vandemoortele et al., 2019), a GFP or RFP-fused construct for live imaging (Dunst and Tomancak, 2019), a biotin ligase-fused construct for proximity labeling (Bosch et al., 2020), or an RFP-GFP tandem fluorescent-fused construct to monitor autophagic degradation in lysosomes (Kimura et al., 2007). Transgenesis methods introducing a tagged construct at a secondary site are well established in *Drosophila* (Venken and Bellen, 2014). Nevertheless, it is still labor-intensive work to generate multiple transgenic strains for each gene of interest.

Autophagy, an intracellular degradation pathway in eukaryotes, is induced by environmental and developmental stimuli (Klionsky et al., 2021). Developmentally programmed autophagy is seen in a variety of Drosophila tissues, including the salivary gland, muscle, and fat body (Fujita et al., 2017; Berry and Baehrecke, 2007; Rusten et al., 2004). It is thought that developmental autophagy allows the degradation and turnover of the cytosolic materials required for tissue remodeling during metamorphosis (Fujita et al., 2017). In the fat body, autolysosomes, a hybrid organelle of autophagosome and lysosome, are enlarged and accumulate massively at the last larval and white prepupal stage (Rusten et al., 2004; Butterworth et al., 1988; Butterworth and Forrest, 1984). The steroid hormone ecdysone, the master regulator of insect development, drives developmental autophagy in the fat body through downregulation of the Akt/PI3K axis (Rusten et al., 2004). However, it is unclear whether the robust induction of autophagy by ecdysone is the most important cause of the accumulation of enlarged autolysosomes. Autophagic flux has not been explored yet; therefore, it is still possible that a delay in autolysosome turnover resulted in the accumulation of enlarged autolysosomes in the last instar larval and prepupal fat body.

Here we have developed a Drosophila toolkit of HA Frankenbody, a genetically encoded single-chain variable fragment for the HA epitope tag (Zhao et al., 2019). Since the FlyORFeome project generates and distributes a comprehensive *in vivo* genome-wide UAS-ORF-3xHA library of Drosophila (Bischof et al., 2012), the HA Frankenbody toolkit is powerful for functional analysis of POI. This tool allowed us to do live-imaging of HA-tagged constructs in Drosophila tissues. By exploiting the pH sensitivity of GFP, GFP-mCh tandem fluorescent-tagged (TF) HA Frankenbody visualized the autophagic degradation of HA-tagged proteins. Additionally, TF-tagged HA Frankenbody revealed a block in autophagic flux in the last instar larval and prepupal fat bodies, probably due to insufficient acidification of lysosomes. Altogether, an HA Frankenbody toolkit enables various protein localization and dynamics studies, including the autophagic degradation assay, by simply crossing HA Frankenbody transgenic flies with HA-tagged POI flies.

## Results

### HA Frankenbody recognizes HA-tagged constructs in live Drosophila tissues

We generated a series of transgenic lines harboring fluorescent protein-fused HA Frankenbody under UAS or QUAS enhancer sequences (Table 1) to express the HA Frankenbody in Drosophila tissues. Binary expression systems allow the expression of genes in a tissue of interest (Brand and Perrimon, 1993; Riabinina et al., 2015). To express both the HA-tagged protein and the HA Frankenbody construct under the control of the UAS sequences, the progenies need three transgenes; 1) a tissue-specific GAL4 driver, 2) UAS-gene of interest (GOI)-HA, and 3) UAS-HA Frankenbody (Fig. 1A). For this purpose, UAS-HA Frankenbody was combined with several GAL4 lines, such as Cg-GAL4, DMef2-GAL4, pnr-GAL4, or da-Gal4 (Table 1 and Fig. 1A). These are drivers for fat bodies, muscle cells, larval epidermal cells, and broad tissues. We used larval epidermal cells (LECs) to assess the system because LECs are mononuclear and form a monolayer suitable to observe intracellular organelles. First, we expressed HA Frankenbody-GFP in the absence of HA-tagged protein. HA Frankenbody-GFP diffusely localized in both cytosol and nucleus, the same as GFP, in the absence of a HA-tagged construct (Fig. 1B; LacZ, control). Next, we co-expressed HA Frankenbody-GFP and HA-tagged organelle markers. *γ*COP, TOM20, and βTub97EF are markers for Golgi, mitochondria, and microtubules, respectively. The localization of HA Frankenbody-GFP changed upon expression of *γ*COP-3xHA, TOM20-3xHA, or βTub97EF-3xHA (Fig. 1B). HA Frankenbody-GFP signal colocalized entirely with the anti-HA immunostaining signal (Fig. 1B, magnified and line-plot). We also confirmed their colocalization in larval body wall muscles (LBWMs) (Fig. S1). These results show that the HA Frankenbody visualizes HA-tagged proteins in Drosophila tissues.

**Table 1.** anti-HA frankenbody flies.

| Category | Chr. | Genotype |
| --- | --- | --- |
| Transgenic lines | 1;2 | w; UAS-Frankenbody<sup>HA</sup>:GFP (attP40) |
|  | 1;2 | w; UAS-Frankenbody<sup>HA</sup>:mCherry (attP40) |
|  | 1;2 | w; UAS-Frankenbody<sup>HA</sup>:GFP:mCherry (attP40) |
|  | 1;3 | w; QUAS-Frankenbody<sup>HA</sup>:GFP (attP2) |
|  | 1;3 | w; QUAS-Frankenbody<sup>HA</sup>:mCherry (attP2) |
|  | 1;3 | w; QUAS-Frankenbody<sup>HA</sup>:GFP:mCherry (attP2) |
| Combined lines | 1;2 | w; Cg-GAL4, UAS-Frankenbody<sup>HA</sup>:GFP |
|  | 1;2 | w; Cg-GAL4, UAS-Frankenbody<sup>HA</sup>:mCherry |
|  | 1;2 | w; Cg-GAL4, UAS-Frankenbody<sup>HA</sup>:GFP:mCherry |
|  | 1;2;3 | w; UAS-Frankenbody<sup>HA</sup>:GFP; pnr-GAL4 |
|  | 1;2;3 | w; UAS-Frankenbody<sup>HA</sup>:mCherry; pnr-GAL4 |
|  | 1;2;3 | w; UAS-Frankenbody<sup>HA</sup>:GFP:mCherry; pnr-GAL4 |
|  | 1;2;3 | w; UAS-Frankenbody<sup>HA</sup>:GFP; DMef2-GAL4 |
|  | 1;2;3 | w; UAS-Frankenbody<sup>HA</sup>:mCherry; DMef2-GAL4 |
|  | 1;2;3 | w; UAS-Frankenbody<sup>HA</sup>:GFP:mCherry; DMef2-GAL4 |
|  | 1;2;3 | w; UAS-Frankenbody<sup>HA</sup>:GFP; da-GAL4 |
|  | 1;2;3 | w; UAS-Frankenbody<sup>HA</sup>:mCherry; da-GAL4 |
|  | 1;2;3 | w; UAS-Frankenbody<sup>HA</sup>:GFP:mCherry; da-GAL4 |

**Figure 1.**
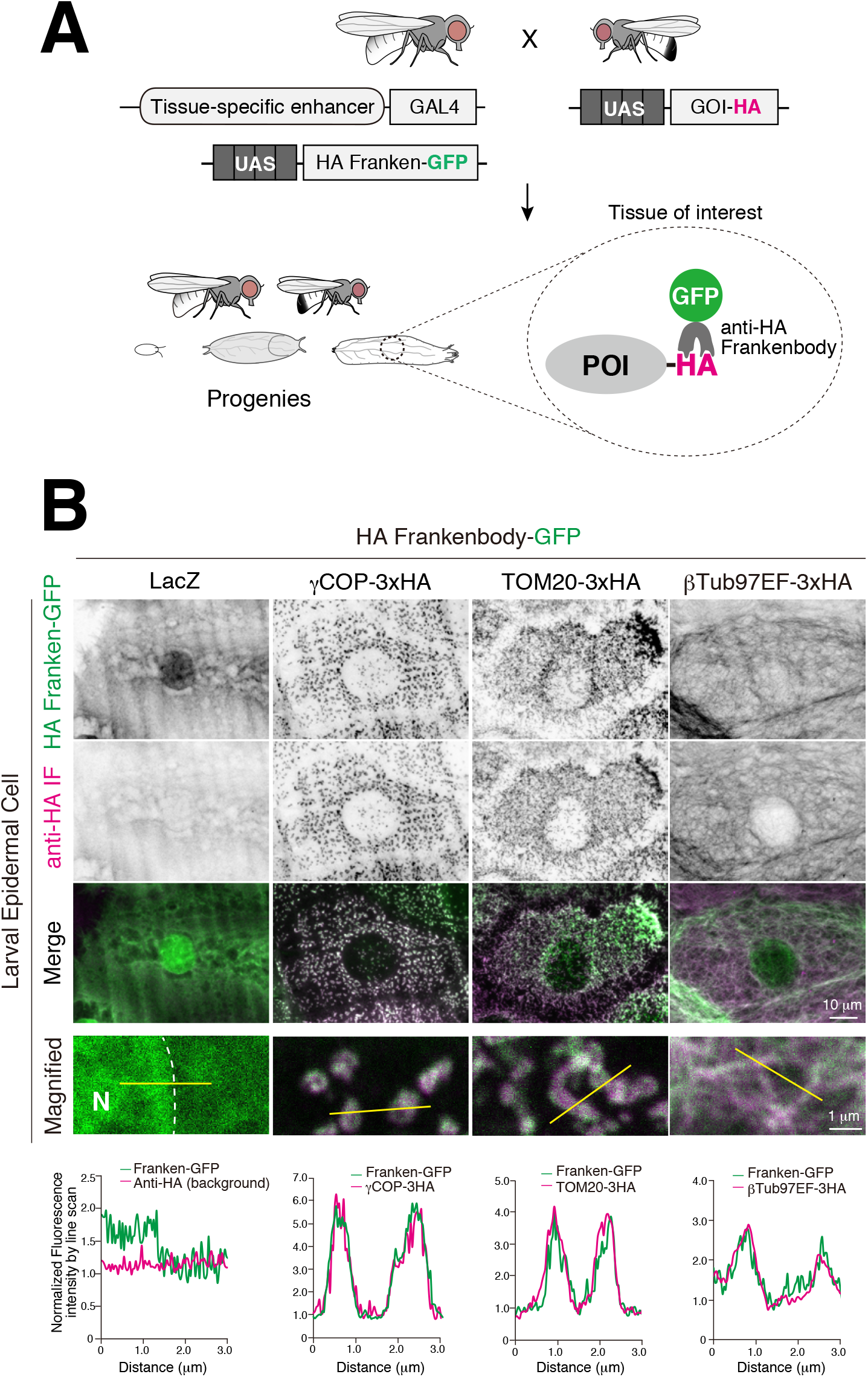
HA Frankenbody recognizes HA-tagged protein in larval epidermal cells. (A) Schematic of the HA Frankenbody strategy using the GAL4-UAS binary expression system. (B) Colocalization of HA Frankenbody-GFP and γCOP-3xHA, TOM20-3xHA, or βTub97EF-3xHA in larval epidermal cells, N; nucleus. Line plots profile the yellow line in each panel.

Next, we observed the HA Frankenbody-GFP in live tissues. HA Frankenbody-GFP and HA-tagged protein were co-expressed in LECs, LBWMs, or fat bodies and observed in live tissues by confocal microscopy (Fig. 2A-C, S2). LECs and LBWMs were observed through the cuticle in intact animals. Conversely, fat bodies were dissected since they were too far from the cuticle for a high magnification objective lens. Upon co-expression of 3xHA-tagged organelle markers, HA Frankenbody-GFP showed the typical localization pattern associated with each organelle (Fig. 2, S2). Collectively, HA Frankenbody allows imaging of HA-tagged constructs in live Drosophila tissues.

**Figure 2.**
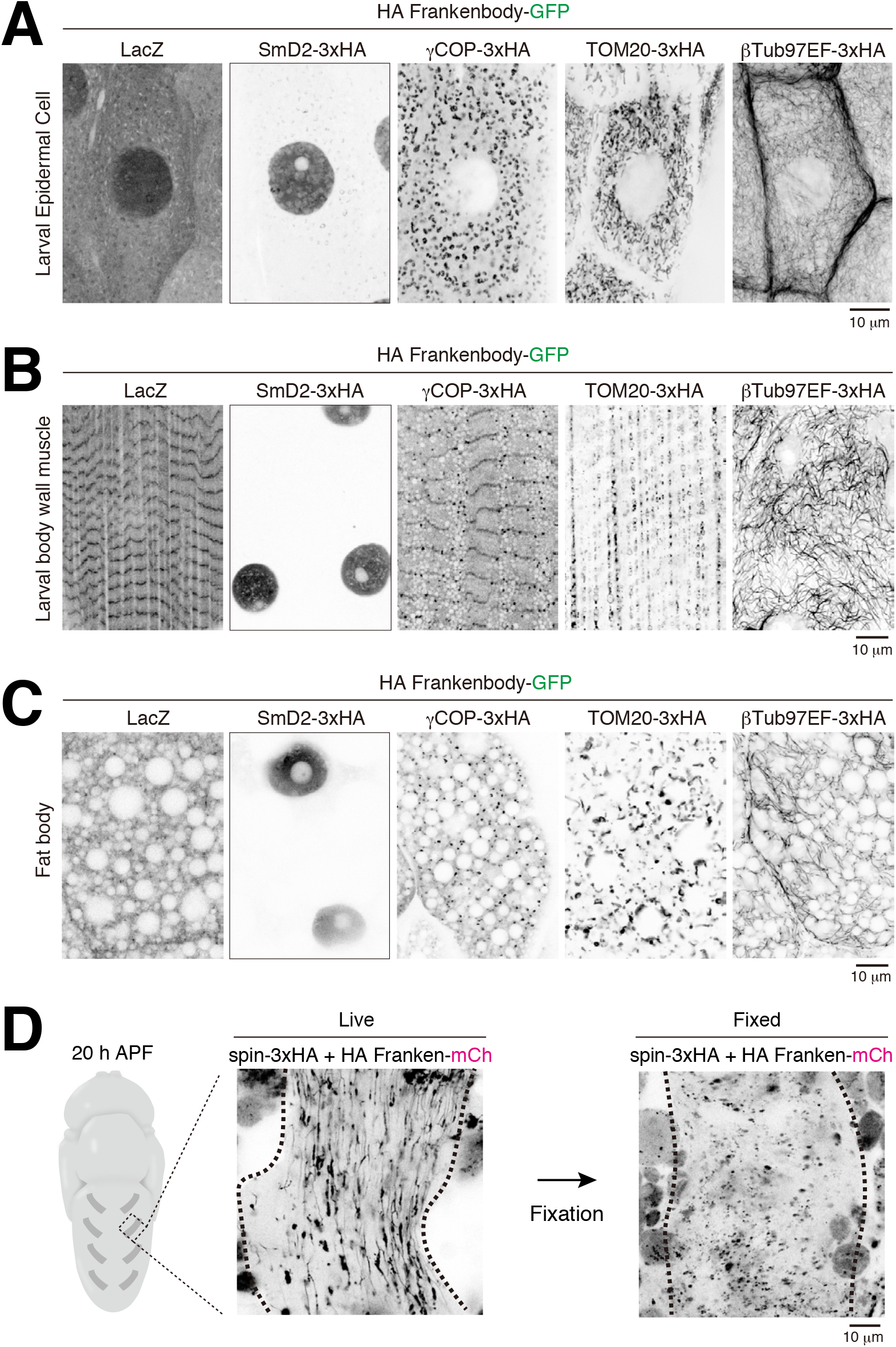
HA Frankenbody allows imaging of HA-tagged constructs in live Drosophila tissues. (A-C) Localization of HA Frankenbody-GFP in cells expressing the indicated 3xHA-tagged constructs in live LECs (A), LBWMs (B), or fat bodies (C). (D) Pupal DIOMs expressing both spinster-3xHA and HA Frankenbody-mCh were observed before (Live) and after (Fixed) fixation.

We recently reported that the spinster-positive tubular autolysosomal (tAL) network functions in the remodeling of the abdominal muscles during metamorphosis (Murakawa et al., 2020). The tAL network is fragile and highly sensitive to dissection and fixation processes (Murakawa et al., 2020; Johnson et al., 2015). Hence, it is almost impossible to observe the tAL network by immunostaining. In contrast to that, HA Frankenbody-mCh visualized the spin-3xHA-positive tAL network in 20 h dorsal internal oblique muscles (DIOMs) (Fig. 2D). Subsequent to dissection and fixation, the structure was lost (Fig. 2D), showing that HA Frankenbody is suitable for imaging of fragile structures that are highly sensitive to fixation.

### TF-tagged HA Frankenbody visualizes the lysosomal degradation of HA-tagged proteins of interest

The tandem fluorescent protein-tagged Atg8 homolog, GFP-mCh-LC3/Atg8, allows the assessment of autophagy flux because of the pH sensitivity of GFP (Kimura et al., 2007). In analogy to this reporter, we generated a transgenic fly carrying UAS-HA Frankenbody-GFP-mCh (Table 1) to monitor the autophagic degradation of HA-tagged protein. Both GFP and mCh fluoresce at the neutral pH of the cytosol (Fig. 3A left). On the other hand, only mCh fluoresces at the acidic pH in lysosomes because the pKa is around 6.0 for GFP and it is quenched in acidic conditions (Fig. 3A right). To validate the TF system, both HA franken-TF and HA-tagged constructs were expressed in the larval fat bodies. Third instar larvae (3IL) were fed or starved for 4 h, and fat bodies were observed (Fig. 3B and C). In fed condition, expression of LacZ, GAPDH1-3xHA, or TOM20-3xHA induced only a low level of mCh-single-positive puncta (Fig. 3B and C). In contrast, expression of ref(2)p-HA, an established autophagy cargo protein (Komatsu et al., 2007; Nezis et al., 2008), induced robust formation of mCh-single-positive puncta, suggesting ref(2)p-HA is degraded in the lysosome even under fed conditions (Fig. 3B and C). We obtained a similar result in 1-week-old adult indirect flight muscles (Fig. S3A, B). Furthermore, nutrient-starvation significantly elevated the lysosomal degradation of the TF-tagged HA Frankenbody in the presence of ref(2)p-HA (Fig. 4B and C), suggesting that the degradation occurs via autophagy. We also noticed that, under fed conditions, ref(2)p-HA expression caused the bright spherical structures positive for both GFP and mCh (Fig. 3B). The size of the GFP-positive spherical structures significantly decreased in parallel with their degradation upon starvation (Fig. 3D-E, S3C-D), suggesting that the spherical structures are liquid or gel-like ref(2)p-droplets (Pircs et al., 2012; Kageyama et al., 2021).

**Figure 3.**
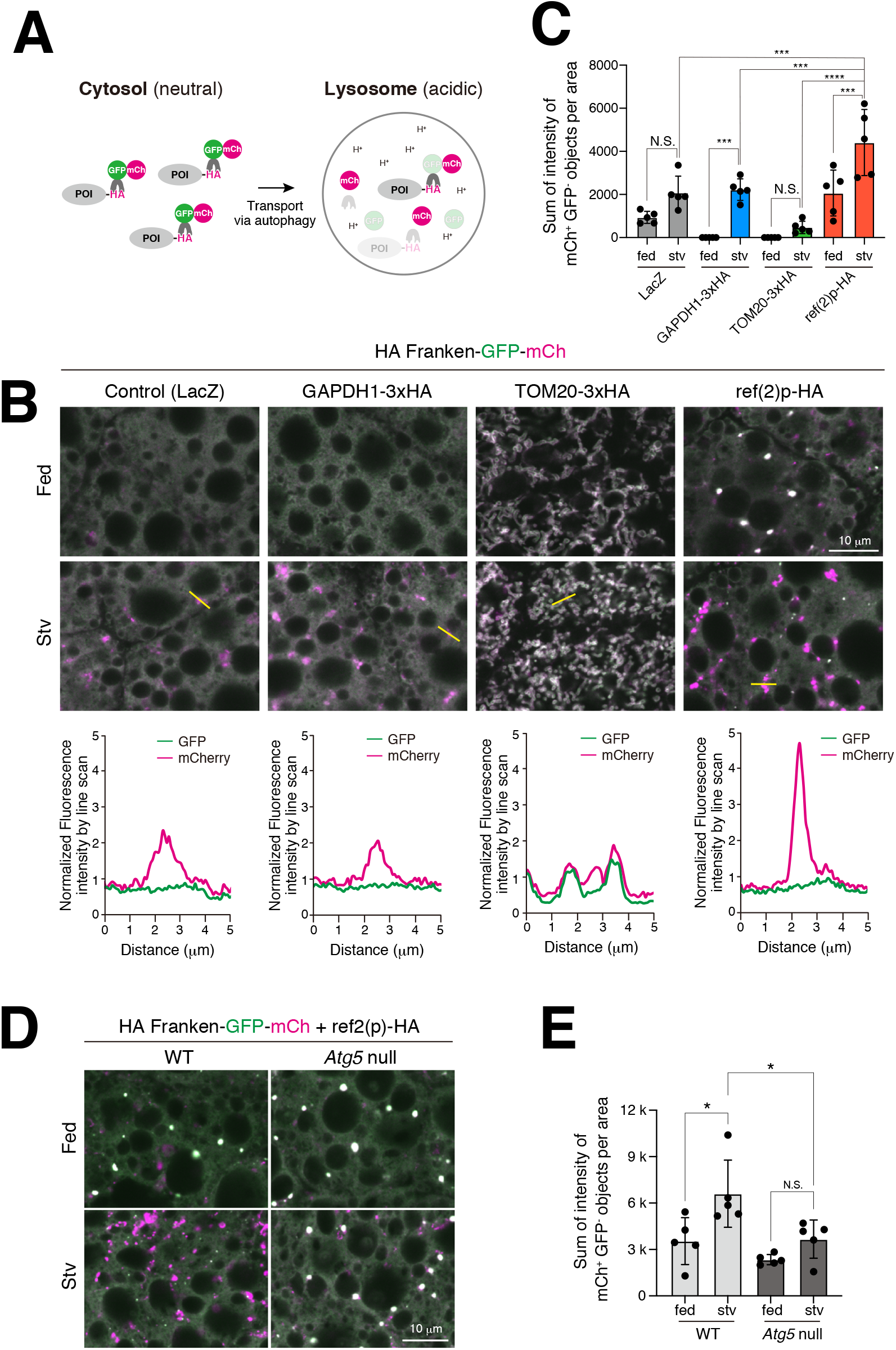
TF-tagged HA Frankenbody visualizes the autophagic degradation of HA-tagged proteins of interest. (A) Schematic of the lysosomal degradation assay using TF-tagged HA Frankenbody. (B and C) Early 3IL larvae expressing HA Frankenbody-GFP-mCh and the indicated HA-tagged construct in fat bodies were fed or starved for 4 h. (B) Confocal images of GFP and mCh channels in each condition. Line plots profile the yellow line in each panel. (C) Quantification of the total intensity of mCh-positive and GFP-negative objects per area; ±SD for 5 images from 5 animals. (D and E) Effect of the loss of *Atg5* on the degradation of HA Frankenbody-GFP-mCh in the fat body co-expressing ref(2)p-HA. (D) Confocal images of GFP and mCh channels in each condition. (E) Quantification of the total intensity of mCh-positive and GFP-negative objects per area; ±SD for 5 images from 5 animals.

**Figure 4.**
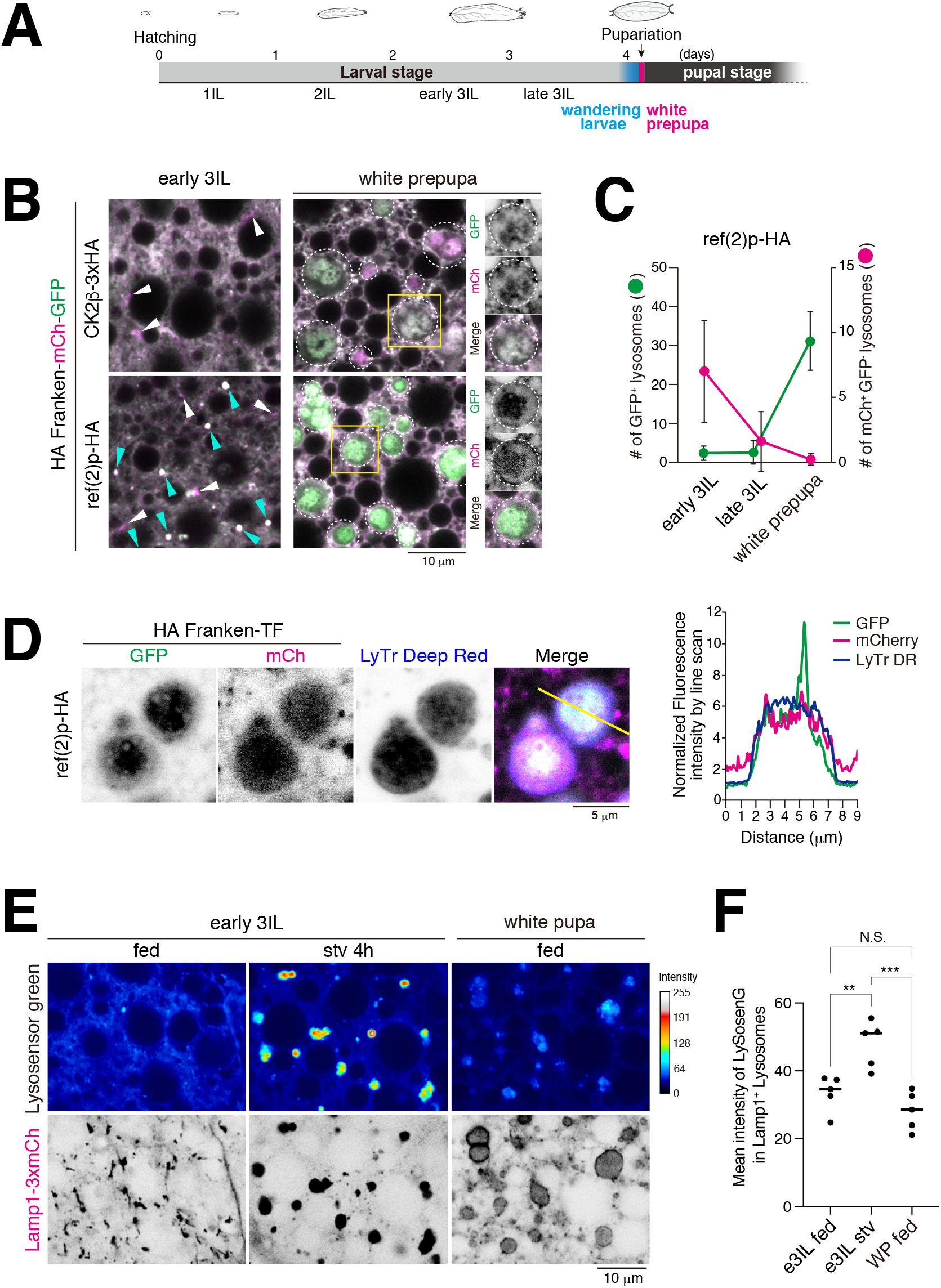
Acidification of lysosomes in the prepupal fat body is regulated. (A) Timeline of fly development from embryo to pupa at 25°C. (B and C) Fat bodies expressing both HA Frankenbody-GFP-mCh and CK2β-3xHA or ref(2)p-HA at the early 3IL or pupariation (white prepupa). (B) Confocal images of GFP and mCh channels under the indicated conditions. mCh-positive and GFP-negative puncta, white arrowhead; both GFP and mCh-positive bright spherical structures, cyan arrowhead; both GFP and mCh-positive enlarged structures, white dotted circle. (C) Number of objects positive for only mCh (blue) or objects positive for both GFP and mCh (pink). (D) Colocalization of GFP, mCh, and LysoTracker Deep Red in white prepupal fat body expressing both HA Frankenbody-GFP-mCh and ref(2)p-HA. Line plots profilethe yellow line in the panel. (E and F) Fat bodies expressing Lamp1-mCh were stained with Lysosensor Green at early 3IL or pupariation (white prepupa). (E) The intensity map shows a representative image of the intensities of LysoSensor Green-positive objects. The median intensity in each image was set as 60. (F) Quantification of mean intensities of Lysosensor Green-positive objects. N=5.

To examine whether the degradation of the TF reporter depends on autophagy, we tested *Atg5* null mutants for the formation of mCh-only puncta. *Atg5* is an essential gene for autophagy (Mizushima et al., 1998; Kim et al., 2016). As shown in Fig. 3 D-E, loss of *Atg5* significantly blocked mCh puncta formation. In addition, the size of GFP-positive spherical structures was not changed by starvation in *Atg5* null mutants (Fig. S3C-D). These results indicate that a complex of ref(2)p-HA and HA Frankenbody-TF was delivered into lysosomes via autophagy. Altogether, the above results demonstrate that the HA Frankenbody-TF visualizes the lysosomal degradation of HA-tagged protein of interest in Drosophila tissues.

### Lysosomal acidification is developmentally regulated in the prepupal fat body

By using time course microscopy to observe fat bodies expressing HA Frankenbody-TF from early 3IL to white prepupa, we found that both GFP and mCh fluoresce in the enlarged structures in the white prepupal fat bodies (Fig. 4A-C, white dotted circle in B). For quantification, the bright spherical structures positive for both GFP and mCh were excluded (Fig. 4B, cyan arrowhead) because they were lysotracker-negative liquid or gel-like ref(2)p-droplets (Fig. S4, 3D-E). Parallel to the increase of GFP-positive objects, the number of mCh-single-positive objects decreased (Fig. 4B-C, white arrowhead). These results suggest that the GFP-positive structures are lysosome-related structures with relatively higher pH. To test this possibility, white prepupal fat bodies expressing both HA franken-TF and ref(2)p-HA were stained with LysoTracker Deep Red, a dye for acidic organelles. As shown in Fig. 4D, the enlarged GFP-positive structures were also positive for LysoTracker dye, indicating that they are acidic lysosomal compartments. Although the LysoTracker probe accumulates and fluoresces in the acidic compartments, its intensity is largely independent of pH (Guha et al., 2014). In contrast, the LysoSensor dye exhibits a change in fluorescent intensity with luminal pH. Therefore, we compared the intensity of Lysosensor Green in the compartments in starved 3IL or white prepupal fat bodies. The LysoSensor showed a significantly lower intensity in the white prepupal fat body than the starved 3IL fat body (Fig. 4E-F), indicating that lysosomal acidification is regulated in the prepupal fat bodies.

### Lysosomal function is developmentally downregulated in an autophagy-independent manner

It is thought that robustly induced autophagy is the dominant cause for enlargement of lysosomal compartments in the last instar larval fat body (Rusten et al., 2004). To test this model, we examined if there was a correlation between autophagy induction and the size of the lysosome. The number of GFP-Atg8 puncta (autophagosomes) and the size of LysoTracker Red-positive compartments (lysosomes) were quantified in starved 3IL or white prepupal fat bodies (Fig. 5A-D) (Kabeya et al., 2000). The ratio of the lysosome area per number of autophagosomes is significantly higher at the white prepupal stage than the starved 3IL stage (Fig. 5D), suggesting that not only autophagy induction but also other mechanisms contribute to the enlargement of lysosomes. Next, we tested the loss of *Atg5* on the size of Lamp1-3xmCh-positive lysosomes to further examine the contribution of autophagy to the enlargement of lysosomes (Hegedus et al., 2016). Strikingly, the enlargement was observed even in *Atg5* null conditions (Fig. 5E-F). The size of the lysosomes was affected only modestly by the loss of *Atg5* in the white prepupal fat body. Next, we performed a flux assay using chloroquine, a well-known lysosomotropic agent which increases lysosomal pH and vacuolation (Zhou et al., 2020). Chloroquine feeding significantly increased the size of the Lamp-positive lysosomes in white prepupal LECs (Fig. 5G-H, Fig. S5C). In sharp contrast, the enlargement by chloroquine was hardly seen in the fat body at the same stage (Fig. 5G-H, Fig. S5D). Collectively, these results show that excess autophagy is not the dominant cause for the enlargement of lysosomes, and lysosomal function is downregulated in the prepupal fat body.

**Figure 5.**
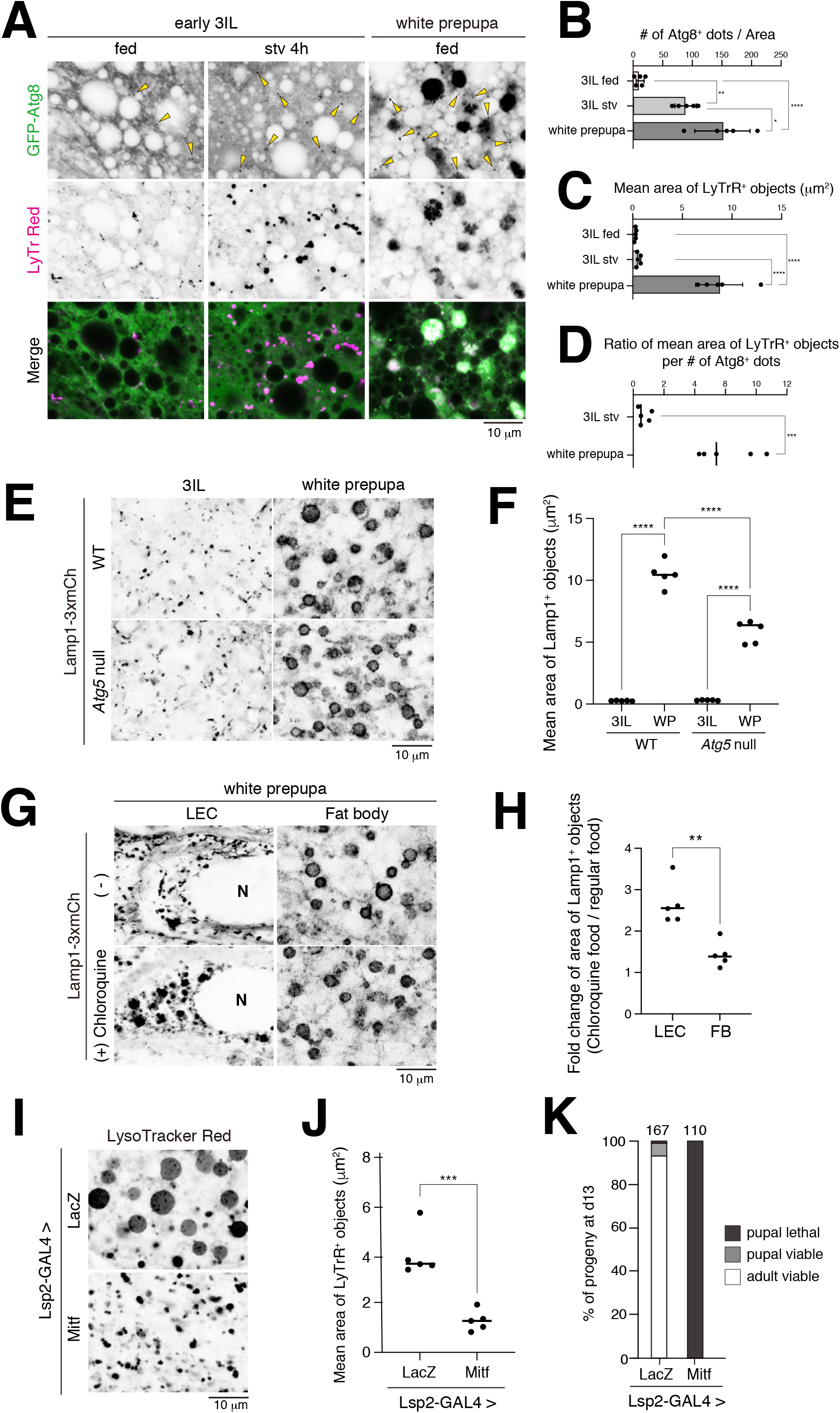
Lysosomes are developmentally downregulated in an autophagy-independent manner. (A-D) Fed or starved fat bodies expressing GFP-Atg8 were stained with LysoTracker red at early 3IL or pupariation (white prepupa). (A) Confocal images of GFP-Atg8 and LysoTracker red under the indicated conditions. GFP-Atg8-positive small puncta, yellow arrowhead. (B) The number of small GFP-Atg8 dots per area (C) Mean area of LysoTracker red-positive objects. (D) The ratio of the area of LysoTracker red-positive objects per number of small GFP-Atg8 dots; ±SD for 5 images from 5 animals. (E and F) The effect of *Atg5* null mutation on the size of lysosomes at early 3IL or pupariation (white prepupa). (E) Confocal images of Lamp1-3xmCh under the indicated conditions. (F) The mean area of Lamp1-3xmCh-positive objects; ±SD for 5 images from 5 animals. (G and H) The effect of chloroquine feeding on the size of lysosomes in LECs or fat bodies at pupariation, N; nucleus. (G) Confocal images of Lamp1-3xmCh under the indicated conditions. (H) Fold change of the area of Lamp1-3xmCh-positive lysosomes upon chloroquine feeding. (I-K) Effect of Mitf overexpression on the size of LysoTracker Red-positive lysosomes in white prepupal fat bodies. (I) Confocal images of LysoTracker Red in white prepupal fat bodies expressing LacZ (control) or Mitf. (J) The mean area of LysoTracker Red-positive objects; ±SD for 5 images from 5 animals. (K) Percentage of viable and lethal progeny at 13 days after egg laying with fat body-targeted Mitf overexpression.

Lastly, we explored the biological significance of the downregulation of lysosomes in the fat body. For this purpose, we tested fat body-targeted Mitf overexpression by Lsp2-GAL4 on the enlargement of lysosomes and the development of Drosophila. Mitf is the single MiTF/TFE family gene in Drosophila and is a master regulator of the autophagy-lysosome pathway (Zhang et al., 2015; Bouché et al., 2016). Thus, Mitf overexpression upregulates lysosomal functions through transcription activation of a series of lysosome-related genes, including v-ATPase subunits required for the acidification of lysosomes (Bouché et al., 2016). Mitf overexpression markedly suppressed the enlargement of lysosomes in the white prepupal fat body (Fig. 5I-J), probably through the forced activation of lysosomes. It is important to note that Lsp2-GAL4 drives expression only after late 3IL. At 13 days after egg laying, almost all progenies became adults in the control (Fig. 5K). By contrast, fat body-targeted Mitf overexpression induced severe lethality at the pupal stage (Fig. 5K), suggesting the downregulation of lysosomes in the last instar larval and prepupal fat body plays a role in the metamorphosis of Drosophila.

## Discussion

A toolkit of HA Frankenbody allows a versatile localization and dynamics analysis of HA-tagged POI in Drosophila. A combination of HA Frankenbody and the Zurich FlyORFeome library enables large-scale localization screening *in vivo*. The FlyORFeome project aims to develop a comprehensive *in vivo* genome-wide UAS-ORF library. Notably, the UAS-transgenes were fused with a 3xHA tag at the C-terminus (Bischof et al., 2012). Figures 1 and 2 demonstrated that our strategy was powerful for *in vivo* subcellular localization analysis. Cytosolic HA Frankenbody does not recognize the HA tag exposed in the luminal or extracellular space. However, we think this feature is advantageous for analyzing the membrane topology of a POI. If an inserted HA tag is recognized by HA Frankenbody, one can tell that the HA-fused segment is exposed to the cytosol.

Immunostaining is a standard protocol for the localization analysis of epitope-tag fused constructs. Nevertheless, chemical fixation and permeabilization by detergents often cause artifacts such as changes in the morphology of organelles, loss of epitopes, or mislocalization of proteins (Richter et al., 2018). As such, we analyzed the tAL network, which is highly sensitive to a normal immunostaining process (Murakawa et al., 2020). A combination of spin-3xHA and HA Frankenbody-mCh succeeded in imaging this tubular network in live muscle cells (Fig. 2D). Hence, the HA Frankenbody system is especially suitable for fragile structures labeled with HA-tagged POI where fluorescent protein-tagging is not a viable option.

TF-tagged HA Frankenbody enables monitoring of autophagic degradation of HA-fused POI in lysosomes (Fig. 3). ref(2)p-HA, a fly homolog of p62/SQSTM1 that is an established autophagy cargo, was more efficiently degraded compared with other constructs in both muscles and fat bodies (Fig. 3B-C). We also confirmed that the degradation of ref(2)p-HA depends on autophagy (Fig. 3D-E). Using a combination of TF-tagged HA Frankenbody and HA-fused FlyORF library allows for the assessment of autophagic degradation of POIs without establishing a new tandem-fluorescent protein-fused transgenic line. Autophagy specifically sequesters target organelles, including ER, mitochondria, peroxisomes, and lysosomes, for degradation (Morishita and Mizushima, 2019); however, the physiological roles of the organelle-phagy are not fully understood, especially in multicellular organisms. Thus, it would be interesting to examine the degradation of markers for each organelle in various tissues, developmental stages, and nutrient conditions by the TF-tagged HA Frankenbody strategy. Also, TF-tagged HA Frankenbody is a powerful tool to explore cargo specificity in autophagy. Most selective autophagy cargoes have a short linear sequence motif, called LIR, LRS, or AIM, that physically interacts with Atg8 family proteins (Kalvari et al., 2014). It is now feasible to test all predicted AIM-containing proteins by combining TF-tagged HA Frankenbody and the FlyORF library. A systematic approach would provide insights into cargo recognition by Atg8 on the autophagic membrane.

We observed accumulated and enlarged autolysosomes in the last instar larval and prepupal fat body, consistent with previous reports (Rusten et al., 2004; Butterworth et al., 1988; Butterworth and Forrest, 1984). However, we propose here a new mechanistic model explaining this phenomenon. Our data support a model where decreased autophagic flux is the dominant cause of autolysosome accumulation. First, lysosomal acidification was regulated in prepupal fat bodies (Fig. 4E-F). Second, blockade of lysosomal activity by chloroquine feeding did not significantly affect the size of lysosomes at this stage (Fig. 5E-F). Third, autophagy was dispensable for the enlargement of lysosomes (Fig. 5G-H), indicating that lysosomal activity is developmentally downregulated. The next question to be addressed is the mechanism of the downregulation of lysosomal activity in the last instar larval and prepupal fat body. Since lysosomal acidification was altered compared to the early 3IL stage (Fig. 4E-F), the expression level of *vha* genes, encoding V-ATPase subunits required for lysosomal acidification, or assembly of V0 and V1 V-ATPase subcomplex, could be attenuated (Collins and Forgac, 2020). The regulation of lysosomal pH is complex, and it could be affected by anions, cations, membrane voltage, lysosomal lipids, and so on (Mindell, 2012; Sillence, 2013). Therefore, it is possible that the expression of other lysosome-related genes, such as TPPML, is developmentally regulated in the fat body (Wong et al., 2012). Alternatively, an elevated basic metabolite could be a cause of the downregulation of lysosomes. Comparative RNA-seq or metabolome analysis of early 3IL and white prepupal fat bodies would give valuable clues to the mechanism behind lysosomal swelling.

What is the biological significance of the block in autophagic flux in fat bodies? Forced activation of lysosomes by Mitf overexpression suppressed the accumulation of enlarged lysosomes in the fat body (Fig. 5I-J). Moreover, Mitf overexpression in the last instar fat body induced severe lethality at the pupal stage (Fig. 5K), although its precise mechanism has not yet been revealed. These results suggest that the downregulation of lysosomes in the fat body has a beneficial role in Drosophila development. We hypothesize that the last instar larvae store autolysosomes in fat bodies as a nutrient source for the next harsh starvation period, the pupal stage. The accumulated autolysosomes in the larval fat body might therefore be degraded and recycled in the pupal stage. Supporting this idea, time course electron microscopy of the fat body showed that the number of autophagic vacuoles peaked at 1 d APF and then decreased during the remainder of metamorphosis (Butterworth et al., 1988).

Alternatively, degradation of endocytosed hemolymph proteins or plasma membrane proteins may have some kind of deleterious effect in the fat body, and the ecdysone hormone blocks their degradation for proper development. The exciting questions to be explored are; 1) What are the dynamics of the enlarged autolysosomes in the pupal fat body? 2) What is the effect of forced blockade of autolysosome turnover throughout the pupal period? and 3) What is the mechanism of how fat body-targeted Mitf overexpression induces pupal lethality?

Besides the utilities described in this paper, HA Frankenbody could be applied to proximity labeling of HA-tagged POI by HA Frankenbody-APEX2 or BioID (Bosch et al., 2021), visualization of HA-POI in the extracellular space *in vivo* by a secreted form of HA Frankenbody, and so on. Our study provides a new platform for the functional analysis of HA-tagged proteins *in vivo*. This strategy will be widely used, contributes to the fly community, and is also being applied to *in vivo* studies in other model organisms, such as worms, amphibians, zebrafish, plants, and mice.

## Materials and methods

### Reagents and antibodies

The following antibodies were used: mouse anti-HA (1:1000; clone M180-3; MBL, Nagoya, Japan), anti-mouse IgG Alexa Fluor 568 conjugate (1:1000; Thermo Fisher Scientific, Waltham, MA), LysoTracker Red DND-99 (Thermo Fisher Scientific, Waltham, MA), LysoTracker Deep Red (Thermo Fisher Scientific, Waltham, MA), Lysosensor Green DND-189 (Thermo Fisher Scientific, Waltham, MA), and Chloroquine diphosphate (TCI, Tokyo, Japan).

### Drosophila strains

Flies were reared at 25°C unless otherwise stated. For tissue-targeted gene expression, the following drivers were used; DMef2-GAL4 for muscle, Cg-GAL4 for the fat body, pnr-GAL4 for larval epidermal cells, and da-GAL4 for broad tissues. UAS*-LacZ* was used as a control. Genotypes are described in Table S1. All genetic combinations were generated by standard crosses. Genotypes of flies used in this study include the following: (1) *Atg5*^*5cc5*^/*FM7 actin-GFP* (from JH. Lee; *Atg5* null), (2) *endo_promoter-Lamp:3xmCherry1-9M* (from G. Juhasz), (3) *pnr-GAL4* (from D. Umetsu), (4) *w; UAS-HA:Ref(2)P_7M/TM3 Sb* (from S. Loh), (5) *y w; UAS-GFP:Atg8a* (from T. Neufeld), (6) *w; UAS-Mitf* (96E) (from F. Pignoni), (7) *w*^*1118*^; *P{w*^*+mC*^*=Cg-GAL4*.*A}2* (Bloomington Drosophila Stock Center, BL7011; *Cg-GAL4*), (8) *y*^*1*^ *w*^*\**^; *P{w*^*+mC*^*=GAL4-Mef2*.*R}3* (BL27390; *DMef2-GAL4*), (9) *w*^*1118*^; *P{w*^*+mC*^*=UAS-lacZ*.*B}Bg4-1-2* (BL1776; *UAS-LacZ*), (10) *y w; P{w[+mC]=Lsp2-GAL4*.*H}3* (BL6357; Lsp2-GAL4), (11) *UAS-Smd2:3xHA* (FlyORF, Zurich ORFeome project, F003874), (12) *UAS-γCOP:3xHA* (F000931), (13) *UAS-βTub97EF:3xHA* (F001206), (14) *UAS-Tom20:3xHA* (F003545), (15) *UAS-Gapdh1:3xHA* (F002305), (16) *UAS-CkIIβ:3xHA* (F004937), (17) *UAS-Pex11:3xHA* (F004080), (18) *UAS-Atg18:3xHA* (F002805), (19) *UAS-Atg5:3xHA* (F003001). New genotypes generated during this study include the following: (20) *w*^*1118*^; *UAS-Spinster:3xHA*^*attP40*^, (21) *w*^*1118*^; *UAS-Frankenbody*^*HA*^:*GFP*^*attP40*^, (22) *w*^*1118*^; *UAS-Frankenbody*^*HA*^:*mCherry* ^*attP40*^, (23) *w*^*1118*^; *UAS-Frankenbody*^*HA*^:*GFP:mCherry* ^*attP40*^,(24) *w*^*1118*^; *QUAS-Frankenbody*^*HA*^:*GFP*^*attP2*^, (25) *w*^*1118*^; *QUAS-Frankenbody*^*HA*^:*mCherry* ^*attP2*^, (26) *w*^*1118*^; *QUAS-Frankenbody*^*HA*^:*GFP:mCherry* ^*attP2*^, (27) *w*^*1118*^; *UASt-HA-Frankenbody:GFP/CyO; pnr-GAL4/TM6c Sb Tb*, (28) *w*^*1118*^; *UASt-HA-Frankenbody:GFP/CyO; DMef2-GAL4/TM6c Sb Tb*, (29) *w*^*1118*^; *Cg-GAL4, UASt-HA-Frankenbody:GFP/CyO*, (30) *w*^*1118*^; *UASt-HA-Frankenbody:mCherry/CyO; DMef2-GAL4/TM6c Sb Tb*, (31) *w*^*1118*^; *Cg-GAL4, UASt-HA-Frankenbody:GFP:mCherry/CyO*.

### Preparation of the fat body

Laid eggs were reared at 21°C on standard cornmeal/yeast/agar media for 21 h. Then they were reared at 25°C until they grew to 3IL or white prepupa. For nutrient starvation, 2IL larvae were transferred to fresh food and kept at 25°C for 24 h. The 3IL were reared in standard media (fed) or 20% Sucrose/PBS solution for 4 h (starved). For chloroquine feeding, laid eggs were grown in standard media with 3 mg/mL of chloroquine diphosphate until the indicated stage.

### DNA engineering and generation of transgenic flies

Standard molecular biology techniques were used to construct plasmid vectors. The DNA polymerase KOD One (TOYOBO, Tokyo, Japan) was used for PCR amplification of all the objective sequences. For the expression vector of spinster-3xHA, the spinster coding sequence was amplified with the following primer set; 5’-CACCGAATTCACCATGTCGCTGAAACACCAGAAGCAATC-3’ and 5’-CTCGAGGGCAATCTGACCGCGGCTGATC-3’ from the cDNA. Amplified sequence was inserted into pENTR-D-TOPO vector with TOPO cloning (Thermo Fisher Scientific, Waltham, MA). The inserted coding sequence was transferred to the pTWH vector with the Gateway protocol using LR Clonase II Enzyme mix (Thermo Fisher Scientific, Waltham, MA). Construction of the expression vectors of C-terminal fluorescent protein-tagged HA Frankenbody under UAS control (pUASt-attB-Frankenbody^HA^-GFP, pUASt-attB-Frankenbody^HA^ -mCherry, and pUASt-attB-Frankenbody^HA^ -GFP-mCherry) were performed as follows: as a template, pCMV-15F11-HA-mEGFP was used (Zhao et al., 2019) and for pUASt-attB-Frankenbody^HA^ - GFP, Frankenbody^HA^ -GFP fragments were amplified with the following primer sets: 5’-ATAGGGAATTGGGAACACCATGGCCGAGGTGAAGCTGG -3’ and 5’-ACCCTCGAGCCGCGGCTACTTGTACAGCTCGTCCATGCC -3’. For pUASt-attB-Frankenbody^HA^ -mCherry, the fragments of Frankenbody^HA^ and mCherry were amplified with the following primer sets: 5’-ATAGGGAATTGGGAACACCATGGCCGAGGTGAAGCTGG -3’ and 5’-ACCGCTTCCACCTCCACCGCTTCCG -3’ for Frankenbody^HA^, and 5’-GGAGGTGGAAGCGGTATGGTGAGCAAGGGCGAGGAGGA -3’ and 5’-TGGACGAGCTGTACAAGTGACCGCGGCTCGAGGGT -3’ for mCherry. For pUASt-attB-Frankenbody^HA^ -GFP-mCherry, the fragments of Frankenbody^HA^ -GFP and mCherry were amplified with the following primer sets: 5’-ATAGGGAATTGGGAACACCATGGCCGAGGTGAAGCTGG -3’ and 5’-GCATGGACGAGCTGTACAAGGGTGGACCG -3’ for Frankenbody^HA^ -GFP, and for 5’-ACAAGGGTGGACCGATGGTGAGCAAGGGCGAGGA -3’ and 5’-ATGGACGAGCTGTACAAGTGACCGCGGCTCGAGGGT -3’ for mCherry. Each set of fragments was assembled with pUASt-attB vector digested by *Eco*RI and *Not*I using Gibson assembly. The construction of the expression vectors of the C-terminal fluorescent-tagged Frankenbody^HA^ under QUAS control (pQUASt-attB-Frankenbody^HA^ -GFP, pQUASt-attB-Frankenbody^HA^ -mCherry, pQUASt-attB-Frankenbody^HA^ -GFP-mCherry) was performed as follows: the fragments of Frankenbody^HA^ -GFP, Frankenbody^HA^ -mCherry, and Frankenbody^HA^ -GFP-mCherry were amplified with the following primer sets: 5’-GTTAGATCTCACCATGGCCGAGGTG-3’ and 5’-CTTCTCGAGCTACTTGTACAGCTCGTCCATGCCGAG -3’ for Frankenbody^HA^ -GFP and 5’-GTTAGATCTCACCATGGCCGAGGTG -3’ and 5’-CTTCTCGAGTCACTTGTACAGCTCGTCCATGCCGCC-3’ for Frankenbody^HA^ -mCherry and 5’-GTTAGATCTCACCATGGCCGAGGTG -3’ and 5’-CTTCTCGAGTCACTTGTACAGCTCGTCCATGCCGCC -3’ for Frankenbody^HA^ -GFP-mCherry. The amplified fragments were purified and digested by *Bgl*II and *Xho*I. The digested DNA fragments were ligated to pQUASt-attB also digested by *Bgl*II and *Xho*I using a Ligation-Convenience kit (NIPPON GENE, Tokyo, Japan). All the resultant vectors were validated by sequencing, and then they were injected into embryos for P-element or phiC31 insertion (Wellgenetics, Taipei, Taiwan).

### Tissue preparation and immunofluorescence in Drosophila

Muscle preparations in larval fillets and pupal abdomens were performed as previously described (Ribeiro et al., 2011). 3IL or pupae were pinned on a sylgard-covered petri dish in dissection buffer (5 mM HEPES, 128 mM NaCl, 2 mM KCl, 4 mM MgCl_2_, 36 mM sucrose, pH 7.2). The animals were pinned flat and fixed at room temp for 20 min. (4% PFA, 50 mM EGTA, PBS). Then, the samples were unpinned and blocked at room temp for 30 min (0.3% bovine serum albumin, 2% goat serum, 0.6% Triton X100, PBS), incubated with primary antibody overnight at 4°C, washed (0.1% Triton PBS), then incubated with Alexa Fluor568-conjugated secondary antibody (Thermo Fisher Scientific, Waltham, MA) for 2 hours at room temp. The immunostained samples were washed and mounted in FluorSave reagent (Merck Millipore, Burlington, MA). LEC preparations in 3IL larvae abdomens were performed as previously described (Tenenbaum and Gavis, 2016). After dissection and fixation, the larval body wall muscles were removed from the fillet using fine forceps. Then, samples were unpinned and blocked at room temp for 30 min (0.1% Triton X100, 0.3% BSA, PBS). Subsequent steps were performed as described above for muscles.

### LysoTracker and LysoSensor staining of live fat body

Staged larvae were dissected in PBS, and a lobe of the fat body was collected. The fat bodies were stained with LysoTracker Red (1:10,000 in PBS for 2 min), Lysotracker Deep Red (1:1,000 in PBS for 2 min), or Lysosensor Green (1:2,000 in PBS for 2 min) at room temperature.

### Confocal fluorescence microscopy

For imaging of live larval epidermal cells (LECs), white prepupae were mounted between slide-glass and cover-glass with a spacer. For imaging of live larval body wall muscles (LBWMs), third instar larvae were immobilized by making a tiny hole on the head with fine forceps in 4% PFA, and mounted between slide-glass and cover-glass with a spacer. Live fat bodies were mounted between slide-glass and cover-glass with PBS. LECs, LBWMs, and fat bodies were observed on a confocal microscope FV3000 (Olympus, Tokyo, Japan) with a 60x silicone/1.30 NA UPlanSApo (Olympus, Tokyo, Japan). The image acquisition software used was Fluoview (Olympus, Tokyo, Japan). The exported images were adjusted and analyzed using ImageJ.

### Image analyses

We analyzed up to 10 images acquired from 5 lobes of the fat body from 5 animals for each data. For all image analysis, the images were cropped to 81.59 × 81.59 μm and were analyzed using ImageJ macro language. Below is the analysis process in brief. In Fig. 3C, 3E, and Fig. S3B, the intensity of mCh-only positive dots was quantified. The fluorescence intensity of the GFP channel in each pixel was subtracted from the corresponding pixel of mCh intensity. The map was binarized and the mean intensity and area of each object was analyzed. In Fig. S3D, the area of bright spherical structures positive for both GFP and mCh was quantified. Both GFP and mCh channel images were binarized. Then, they were multiplied together to extract a double positive area. The resultant map was used to analyze the area of each object. In Fig. 4C, the number of mCh-positive and GFP-negative acidic compartments was counted using the same procedure as in Fig. 3C and 3E. Both GFP and mCh-positive large objects were manually counted because it was challenging to segment them automatically. In Fig. 4F, the mean intensity of Lysosensor Green of lamp-3xmCh-positive objects was quantified. First, mCh channel was binarized and filtered to exclude noise by size and circularity. Then, the mean intensity of Lysosensor Green was calculated for each binarized mCh-positive object. In Fig. 5B, the number of GFP-Atg8-positive puncta was counted. The GFP channel image was binarized and filtrated to remove noise.

Then, the number of small puncta was counted. To measure the size of lysosomes in the 3IL fat body, the images were binarized and filtrated to remove noise. Then, the area of each object was quantified. In the white prepupal fat body, lysosome size was analyzed manually using Fiji, because of the same issues with segmentation described above.

### Statistics

Each experiment was performed at least 3 times and at least 5 different cohorts of unique flies were analyzed. All replicate experiments were performed in parallel with controls. The SD was used as error bars for bar charts from the mean value of the data. When more than two genotypes or treatments were used in an experiment, the statistical analysis was performed using Tukey’s test or Dunnett’s test on Prism8 and Prism9 software. A 2-tailed unpaired student’s t-test was used to compare two means. p<0.05 was regarded as statistically significant. p<0.05 is indicated with single asterisks, and p<0.001, p<0.0001, or p<0.00001 is marked with double, triple, or quadruple asterisks, respectively.

## Abbreviations used

APF: after puparium formation
ATG: autophagy-related
mCh: monomeric Cherry
DIOM: dorsal internal oblique muscle
GOI: gene of interest
LBWM: larval body wall muscle
LEC: larval epidermal cell
tAL: tubular autolysosome
TF: GFP-mCh tandem fluorescent
POI: protein of interest
V-ATPase: vacuolar H^+^ ATPase
3IL: third instar larvae

## Acknowledgments

We are grateful to F. Pignoni (Upstate Medical Univ.), S. Loh (Univ. of Cambridge), G. Juhasz (Hungarian Academy of Sciences), G. Davis (UCSF), JH. Lee (Univ. of Michigan), D. Umetsu (Tohoku Univ.), T. Neufeld (Univ. of Minnesota), Bloomington Drosophila Stock Center, VDRC, and FlyORF for reagents. We are grateful to J. Mathieu (College de France) for the helpful discussion. We thank the Biomaterials Analysis Division of the Tokyo Institute of Technology for the DNA sequencing. We are grateful to M. Landekic (McGill Univ.) for English editing. This work was supported in part by Grant-in-Aid for Scientific Research (B) from the MEXT (grant number 21H02473, NF), Grant-in-Aid for Scientific Research on Innovative Areas (grant number 20H05315, NF; 18H05527, HK), Japan Science and Technology Agency (JST) PRESTO (grant number JPMJPR18H8, NF), AMED (grant number JP19ek0109285h0003, NF), and the research grant of Astellas Foundation for Research on Metabolic Disorders (NF).

## Author contributions

NF and TM designed the research; TM, TN, KK, FM, and NF performed the experiments; TM and NF analyzed and interpreted data; NZ, TJS, and HK contributed to the materials; NF took the lead in writing the manuscript. All authors provided critical feedback and helped shape the research and manuscript.

## Supplemental figure legends

**Figure S1 HA Frankenbody recognizes HA-tagged protein in the larval body wall muscles**

Colocalization of HA Frankenbody-GFP and anti-HA immunostaining of LacZ, Smd2-3xHA γCOP-3xHA, or TOM20-3xHA in third instar larval body wall muscles.

**Figure S2 HA Frankenbody allows imaging of HA-tagged constructs in live LECs** Localization of HA Frankenbody-GFP in cells expressing a series of 3xHA-tagged proteins in live LECs.

**Figure S3 Autophagic degradation of HA-tagged constructs in indirect flight muscles** (A and B) Indirect flight muscles expressing HA Frankenbody-GFP-mCh and HA-tagged indicated constructs at 1-week-old adult. (A) Confocal images of GFP and mCh channels in the indicated conditions (upper row). Line plots profile the yellow line in each panel (lower row). (B) Quantification of the total intensity of mCh-positive and GFP-negative objects per area; ±SD for 5 images from 5 animals. (C and D) Effect of the loss of *Atg5* on the ref(2)p droplets in the fat body co-expressing ref(2)p-HA and HA Frankenbody-GFP-mCh. (C) Confocal images of GFP channel in the indicated conditions. (D) Mean area of GFP-positive dots; ±SD for 5 images from 5 animals.

**Figure S4 The spherical ref(2)p droplets are not acidic compartments**

Fed 3IL fat bodies expressing both HA Frankenbody-GFP-mCh and ref(2)p-HA were stained with LysoTracker Deep Red. White dotted circles show HA Frankenbody-GFP-mCh-positive ref(2)p droplets.

**Figure S5 Autophagy flux and the size of lysosomes**

(A and B) The effect of *Atg5* null mutation on the size of lysosomes in fed or starved fat bodies at 3IL. (A) Confocal images of Lamp1-3xmCh-positive lysosomes in the indicated conditions. (B) The mean area of Lamp1-3xmCh-positive lysosomes; ±SD for 5 images from 5 animals. (C and D) The effect of chloroquine feeding on the size of lysosomes in LECs or fat bodies at pupariation. (C) Mean area of Lamp1-3xmCh-positive lysosomes in white prepupal LECs raised in regular or chloroquine food. (D) Mean area of Lamp1-3xmCh-positive lysosomes in white prepupal fat bodies raised in regular or chloroquine food.

