## Supplementary figures and images for "A Drosophila Toolkit for Imaging of HA-tagged Proteins Unveiled a Block in Autophagy Flux in the Last Instar Larval Fat Body"

### Supplemental figures

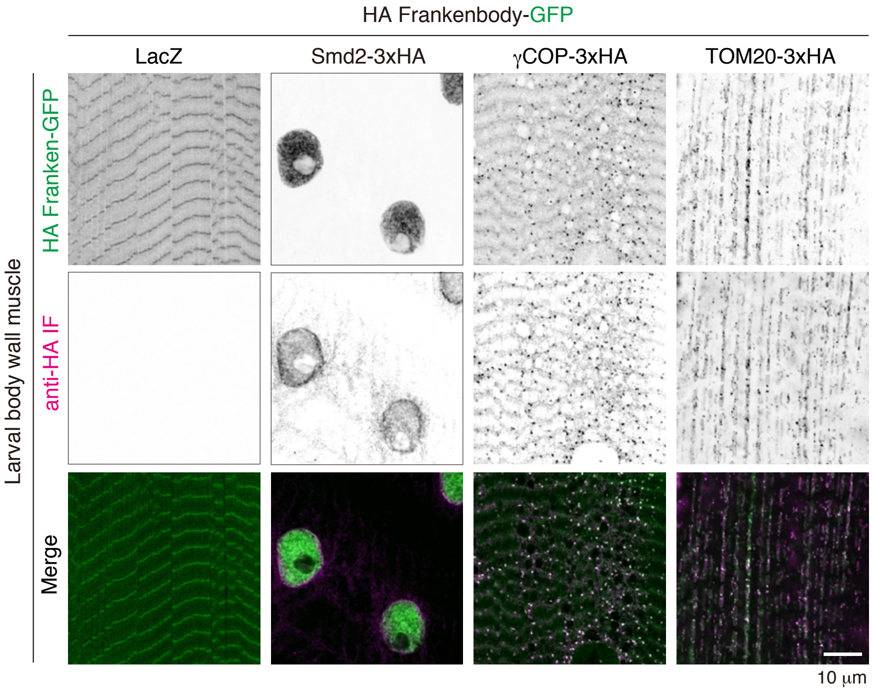

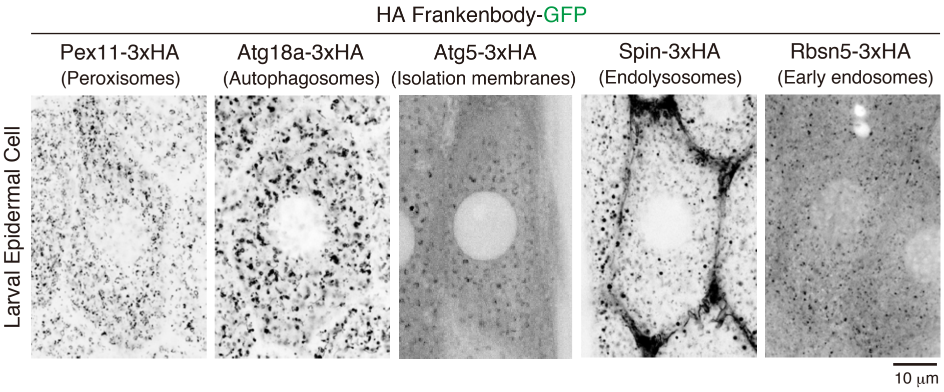

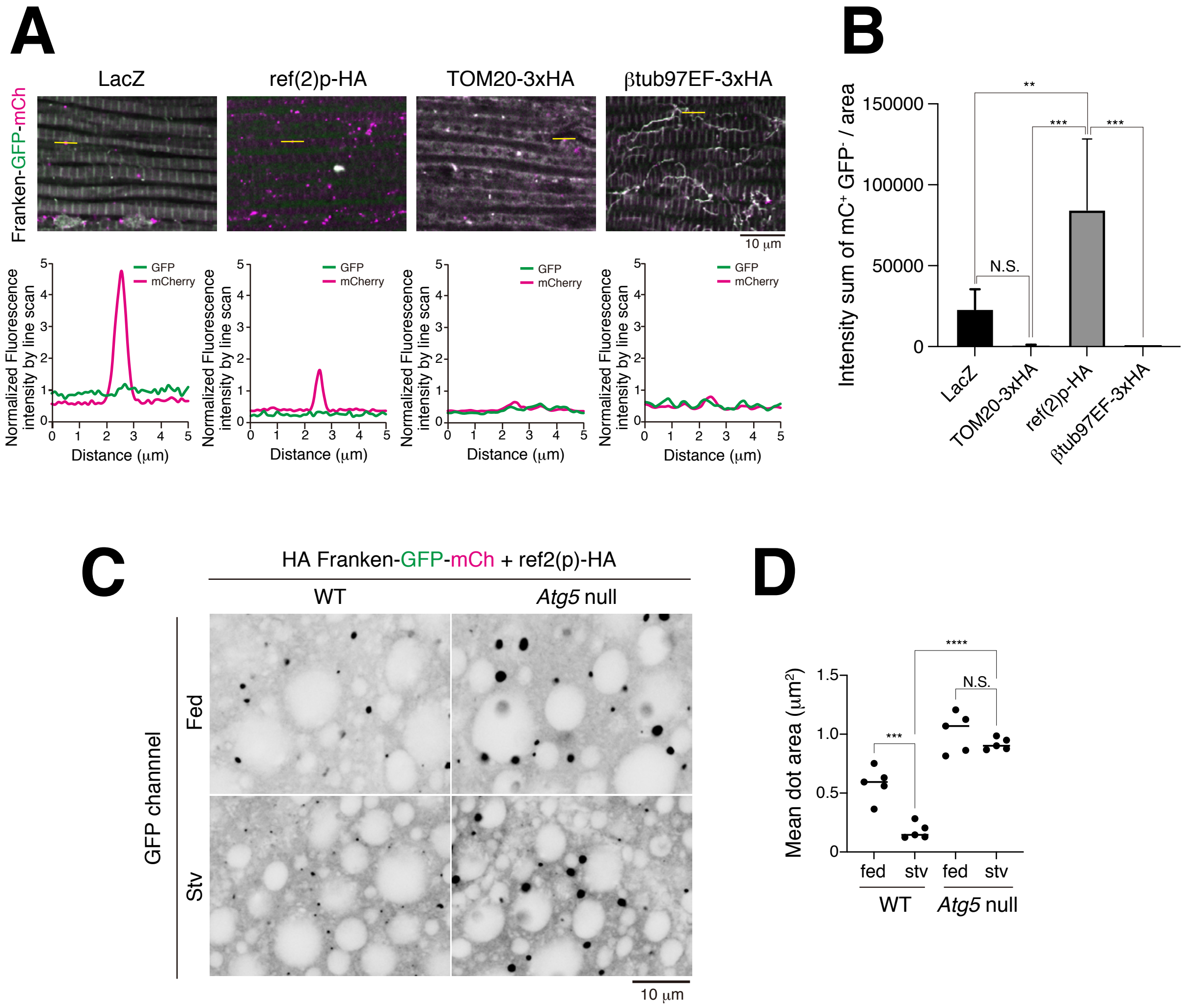

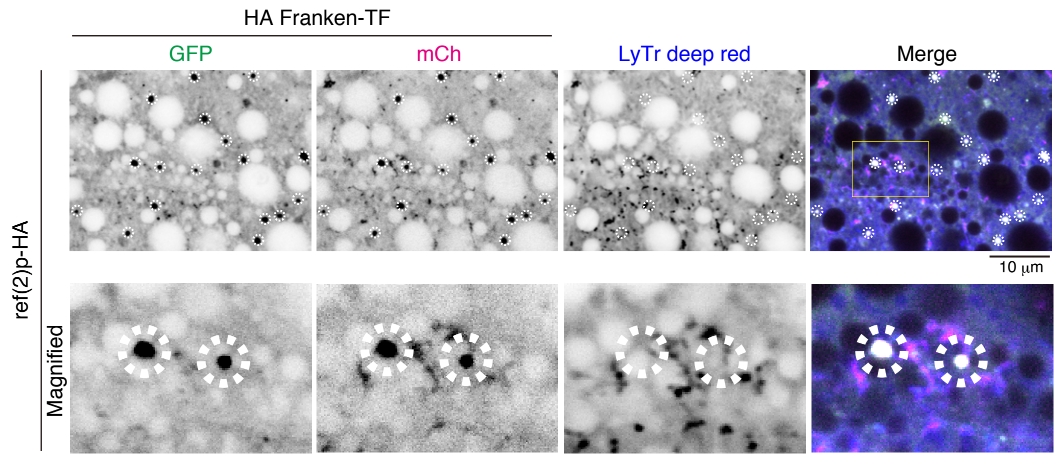

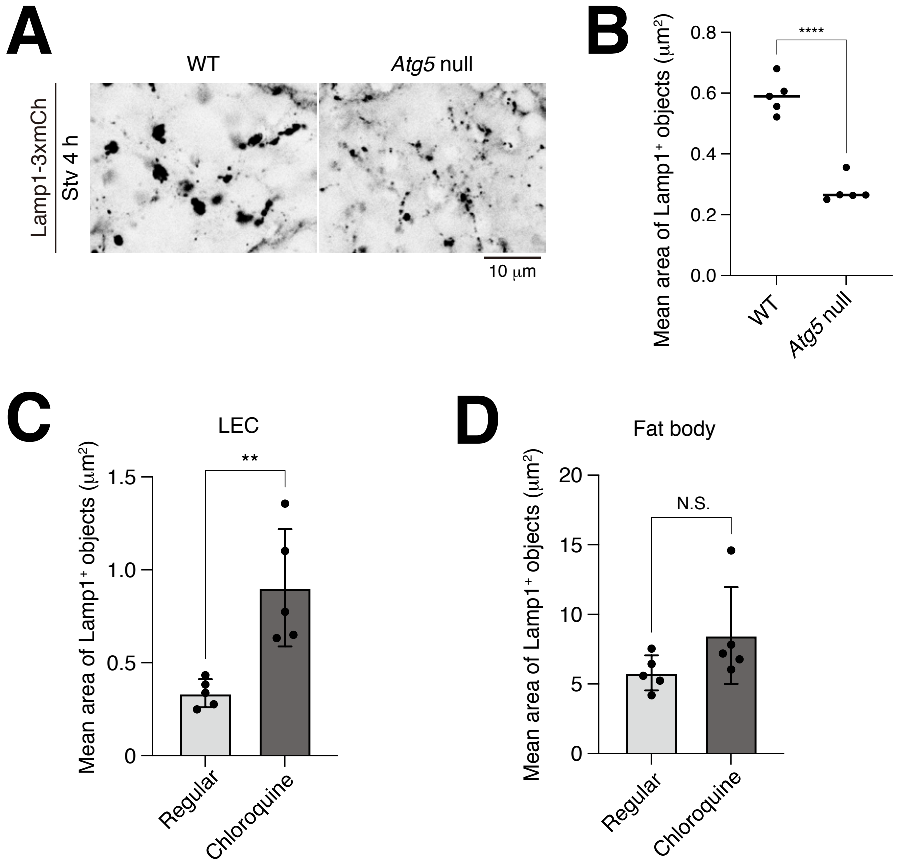
