## Supplemental table for "A Drosophila Toolkit for Imaging of HA-tagged Proteins Unveiled a Block in Autophagy Flux in the Last Instar Larval Fat Body"

### Table S1. Detailed *Drosophila* genotypes shown in figures

| | Panel # | Genotype | Stage | Temp (˚C) |
|---|---|---|---|---|
| Figure 1 | 1B | *UAS-Franken:GFP/UAS-LacZ; pnr-GAL4/+* | 3IL | 25˚C |
| | | *UAS-Franken:GFP/+; pnr-GAL4/UAS-$\gamma$COP:3xHA* | 3IL | 25˚C |
| | | *UAS-Franken:GFP/+; pnr-GAL4/UAS-Tom20:3xHA* | 3IL | 25˚C |
| | | *UAS-Franken:GFP/+; pnr-GAL4/UAS-$\beta$Tub97EF:3xHA* | 3IL | 25˚C |
| Figure 2 | 2A | *UAS-Franken:GFP/UAS-LacZ; pnr-GAL4/+* | white prepupa | 25˚C |
| | | *UAS-Franken:GFP/+; pnr-GAL4/UAS-SmD2:3xHA* | white prepupa | 25˚C |
| | | *UAS-Franken:GFP/+; pnr-GAL4/UAS-$\gamma$COP:3xHA* | white prepupa | 25˚C |
| | | *UAS-Franken:GFP/+; pnr-GAL4/UAS-Tom20:3xHA* | white prepupa | 25˚C |
| | | *UAS-Franken:GFP/+; pnr-GAL4/UAS-bTub97EF:3xHA* | white prepupa | 25˚C |
| | 2B | *UAS-Franken:GFP/UAS-LacZ; DMef2-GAL4/+* | 3IL | 25˚C |
| | | *UAS-Franken:GFP/+; DMef2-GAL4/UAS-SmD2:3xHA* | 3IL | 25˚C |
| | | *UAS-Franken:GFP/+; DMef2-GAL4/UAS-$\gamma$COP:3xHA* | 3IL | 25˚C |
| | | *UAS-Franken:GFP/+; DMef2-GAL4/UAS-Tom20:3xHA* | 3IL | 25˚C |
| | | *UAS-Franken:GFP/+; DMef2-GAL4/UAS-$\beta$Tub97EF:3xHA* | 3IL | 25˚C |
| | 2C | *UAS-Franken:GFP, Cg-GAL4/UAS-LacZ; +/+* | 3IL | 25˚C |
| | | *UAS-Franken:GFP, Cg-GAL4/+; +/UAS-SmD2:3xHA* | 3IL | 25˚C |
| | | *UAS-Franken:GFP, Cg-GAL4/+; +/UAS-$\gamma$COP:3xHA* | 3IL | 25˚C |
| | | *UAS-Franken:GFP, Cg-GAL4/+; +/UAS-Tom20:3xHA* | 3IL | 25˚C |
| | | *UAS-Franken:GFP, Cg-GAL4/+; +/UAS-$\beta$Tub97EF:3xHA* | 3IL | 25˚C |
| | 2D | *UAS-Franken:mCh/UAS-Spinster:3xHA; DMef2-GAL4/+* | 20 h APF | 18℃ |
| Figure 3 | 3B | *UAS-Franken:GFP:mCh, Cg-GAL4/UAS-LacZ; +/+* | early 3IL | 25˚C |
| | | *UAS-Franken:GFP:mCh, Cg-GAL4/+; +/UAS-GAPDH1:3xHA* | early 3IL | 25˚C |
| | | *UAS-Franken:GFP:mCh, Cg-GAL4/+; +/UAS-TOM20:3xHA* | early 3IL | 25˚C |
| | | *UAS-Franken:GFP:mCh, Cg-GAL4/+; +/UAS-ref(2)p:HA* | early 3IL | 25˚C |
| | 3D | *$w^{1118}$/Y; UAS-Franken:GFP:mCh, Cg-GAL4/+; UAS-ref(2)p:HA/+* | early 3IL | 25˚C |
| | | *$Atg5^{5cc5}$/Y; UAS-Franken:GFP:mCh, Cg-GAL4/+; UAS-ref(2)p:HA/+* | early 3IL | 25˚C |
| Figure 4 | 4B | *UAS-Franken:GFP:mCh, Cg-GAL4/UAS-CK2$\beta$:3xHA; +/+* | early 3IL, white prepupa | 25˚C |
| | | *UAS-Franken:GFP:mCh, Cg-GAL4/+; +/UAS-ref(2)p:HA* | early 3IL, white prepupa | 25˚C |
| | 4D | *UAS-Franken:GFP:mCh, Cg-GAL4/+; +/UAS-ref(2)p:HA* | white prepupa | 25˚C |
| | 4E | *endo_promoter-Lamp1:3xmCherry1-9M /CyO* | early 3IL, white prepupa | 25˚C |
| Figure 5 | 5A | *w; Cg-GAL4/UAS-GFP:Atg8; +/+* | early 3IL, white prepupa | 25˚C |
| | 5E | *$w^{1118}$/Y; +/endo_promoter-Lamp1:3xmCherry1-9M; ref(2)p:HA/+* | early 3IL, white prepupa | 25˚C |
| | | *$Atg5^{5cc5}$/Y; +/endo_promoter-Lamp1:3xmCherry1-9M; ref(2)p:HA/+* | early 3IL, white prepupa | 25˚C |
| | 5G | *endo_promoter-Lamp1:3xmCherry1-9M /CyO* | white prepupa | 25˚C |
| | 5I | *UAS-Mitf/+; Lsp2-GAL4/+* | white prepupa | 25˚C |
| Figure S1 | | *UAS-Franken:GFP/UAS-LacZ; DMef2-GAL4/+* | 3IL | 25˚C |
| | | *UAS-Franken:GFP/+; DMef2-GAL4/UAS-SmD2:3xHA* | 3IL | 25˚C |
| | | *UAS-Franken:GFP/+; DMef2-GAL4/UAS-$\gamma$COP:3xHA* | 3IL | 25˚C |
| | | *UAS-Franken:GFP/+; DMef2-GAL4/UAS-Tom20:3xHA* | 3IL | 25˚C |

| | | | | |
|---|---|---|---|---|
| Figure S2 | | *UAS-Franken:GFP/+; pnr-GAL4/UAS-Pex11:3xHA* | white prepupa | 25˚C |
| | | *UAS-Franken:GFP/+; pnr-GAL4/UAS-Atg18a:3xHA* | white prepupa | 25˚C |
| | | *UAS-Franken:GFP/+; pnr-GAL4/UAS-Atg5:3xHA* | white prepupa | 25˚C |
| | | *UAS-Franken:GFP/+; pnr-GAL4/UAS-Spin:3xHA* | white prepupa | 25˚C |
| | | *UAS-Franken:GFP/+; pnr-GAL4/UAS-bTub97EF:3xHA* | white prepupa | 25˚C |
| Figure S3 | S3A | *w; UAS-Franken:GFP:mCh/UAS-LacZ; , DMef2-GAL4/+* | 1-week-old adult | 25˚C |
| | | *w; UAS-Franken:GFP:mCh/+; DMef2-GAL4/UAS-ref(2)p:HA* | 1-week-old adult | 25˚C |
| | | *w; UAS-Franken:GFP:mCh/+; DMef2-GAL4/UAS-TOM20:3xHA* | 1-week-old adult | 25˚C |
| | | *w; UAS-Franken:GFP:mCh/+; DMef2-GAL4/UAS-$\beta$ tub97EF:3xHA* | 1-week-old adult | 25˚C |
| | S3C | *$w^{1118}$/Y; UAS-Franken:GFP:mCh, Cg-GAL4/+; UAS-ref(2)p:HA/+* | early 3IL | 25˚C |
| | | *$Atg5^{5cc5}$/Y; UAS-Franken:GFP:mCh, Cg-GAL4/+; UAS-ref(2)p:HA/+* | early 3IL | 25˚C |
| Figure S4 | | *UAS-Franken:GFP:mCh, Cg-GAL4/+; +/UAS-ref(2)p:HA* | early 3IL | 25˚C |
| Figure S5 | S5A | *UAS-Franken:GFP:mCh, Cg-GAL4/+; +/UAS-ref(2)p:HA* | 3IL | 25˚C |
